# The impact of microplastics on the mice gut microbiome: a meta-analysis

**DOI:** 10.1101/2025.09.22.677931

**Authors:** Anika Kalra, Davide Dominoni, Jelle Boonekamp

**Affiliations:** School of Biodiversity, One Health & Veterinary Medicine, Graham Kerr Building, University of Glasgow, Glasgow, UK

**Keywords:** microplastics, nanoplastics, gut microbiome, Shannon index, relative abundance

## Abstract

**Background:** Microplastics are nearly ubiquitous in the natural environment. There is increasing awareness of the ingestion of microplastics in a wide range of wild animals and humans, but their health and fitness impacts remain poorly understood. Mounting evidence suggests that microplastics could accumulate in the gut, affecting the microbiome and hindering its function. This may constitute a direct pathway through which exposure to microplastics could affect health and disease. However, prior studies on the effects of microplastic exposure on the microbiome show contrasting findings raising the question whether microplastics truly affect the microbiome. Here, we performed the first meta-analysis on the effects of dietary microplastic exposure on the gut microbiome of mice.

**Results:** Overall, we found support for a significant impact of microplastic exposure on the relative abundance of gut bacteria. More in-depth analysis revealed microplastic exposure significantly increased the relative abundance of *Bacillota* and *Pseudomonadota*. In the gut microbiome, *Bacillota* is associated with carbohydrate metabolism and *Pseudomonadota* is associated with facultative anaerobes, thus these changes may have functional consequences.

**Conclusions:** Our findings highlight the importance of distinguishing among differing bacterial phyla when assessing the impacts of microplastics on the gut microbiome as the direction and size of the impact varies by phyla.

## Background

Mass produced plastics have gradually accumulated since the 1950’s and are now considered globally ubiquitous in the natural environment [1–8]. Plastic products break down from natural causes and form “microplastics”, defined by most literature as less than 5 mm, which subsequently break down into “nanoplastics” classed as less than 1 µm [3, 9, 10]. Micro- and nano-plastic particles (henceforth called microplastics) have been found in human food sources such as alcohol, coffee, sugar, honey, salt, bottled water, and tap water, as well as chicken and sheep faeces [6, 11–16]. Microplastics have also been widely reported in marine environments and have been detected in the soft tissue of mussels and oysters [16]. There is growing concern for the health [17] and reproductive [17, 18] implications of plastic ingestion in wild animals, livestock, and humans, as the negative effects of exposure to microplastics are becoming increasingly apparent, but the underlying mechanisms remain unclear.

The main route of microplastics exposure for mammals and humans occurs through ingestion, resulting in the accumulation of microscopic plastic particles in the gastrointestinal tract [4, 6]. It has been hypothesized that the negative health impacts of microplastics exposure could occur through their effects on the gut microbiome composition underpinning chronic disease through metabolic dysfunction, oxidative stress, or inflammation [3–7, 10, 19, 20]. Previous experimental studies on the toxicological impacts of microplastics on gut microbiome parameters have almost exclusively focused on laboratory mice and rat systems [16, 21]. Several of these studies support that experimental microplastics exposure, for example through the diet, results in the accumulation of microplastics in the gut [6], liver and kidney [22], and can induce intestinal microbial dysbiosis through changes in the relative abundance of gut microbiome bacteria and species diversity [5, 22–24]. However, other studies reported no effects of microplastics exposure on the microbiome [21, 25] or different types of changes indicative of dysbiosis [6]. Han et al., 2021 suggests the lack of significant biological and histopathological effects of microplastics exposure found may have been due to experimental differences in dosage and exposure duration. We conclude therefore that it remains unclear whether microplastics affect the gut microbiome and if such effects depend on the dosage and particle size of the microplastics that were used.

Here, we conduct the first meta-analysis on the impacts of microplastics’ exposure on the mice gut microbiome, and whether such effects depend on dosage and particle size. We confined our meta-analysis to experimental laboratory mice studies as they are by far the most common model of studying the toxicological impact of microplastics and are the main model for translation to human diseases [16, 21]. We hypothesize that if microplastics exposure would disrupt the gut microbiome dynamics, then this should lead to an interspecific competition imbalance [8, 26, 27]. If so, then certain bacteria will be outcompeted whilst others thrive, resulting in a reduced microbiome diversity. We also hypothesize increased relative abundance of competitive gut microbiome bacteria and we anticipate these effects to be exacerbated by higher plastic dosages as well as smaller plastic sizes, because smaller particles might penetrate more easily into the epidermal tissues [22, 28].

## Methods

### Data collection

We performed a literature search on the Web of Science and Scopus ending on May 4^th^, 2024 and May 16^th^, 2024, respectively. Our original aim was to capture all research papers on the effects of experimental microplastic exposure on the gut microbiome in terrestrial vertebrates, but during the screening of the literature, it became apparent that the vast majority of suitable papers were on laboratory mice, likely because these are highly suitable systems amenable to experimental exposure to microplastics. Hence, in chronological order, our literature search was conducted as follows: we used the search strings: “microplastics NOT fish NOT invertebrate NOT (mollusc OR arthropod OR worm OR cnidarian OR echinoderm OR sponge) NOT marine NOT aquatic NOT soil AND (microbiome OR microbe)” and “microplastics AND (bird OR mammal OR reptile OR amphibian) AND (microbiome OR microbe)”. This resulted in a total of 405 potentially suitable papers after duplicates across Web of Science and Scopus were removed (Fig. 8). Subsequently, we screened the titles and abstracts for relevancy followed by an in-depth screening of the relevant papers. We adopted the PECO (Population/Exposure/Comparison/Outcomes) format as a framework for applying the inclusion criteria [29]. The exclusion criteria included non-terrestrial species, aquatic, marine or soil environments, engineered treatments or patents, “plasticity” of gut microbiome, introduction of any other independent variables besides microplastic exposure, and microplastic exposure impact on variables outside of the gut microbiome. The first round of screening resulted in 94 papers. At this point, the research question was refined to focus only on mice, resulting in 49 papers (Fig. 8). Out of the 45 papers that were removed in this refinement, only 3 papers (2 studies on chicken and 1 study on toads) would have been suitable considering the original inclusion and exclusion criteria. For the second round of screening, we read the full paper in detail to confirm relevancy and suitability regarding the published data and study design defined as experimental mice studies where there was at least one control group, one plastic particle only treatment group, and microbiome analysis. The second screening round resulted in 38 papers. Finally, if mean and standard deviation data for relative abundance per gut bacteria phyla and/or Shannon’s index was not retrievable, those papers were excluded, resulting in 19 papers ultimately included in our meta-analysis (Fig. 8). See Supplementary Table 1, Additional File 1 for the extracted data by paper ultimately used in our meta-analysis (Table S1).

Gut microbiome dynamics are often expressed in terms of relative abundance for each taxon included in the study, and taxa alpha diversity indices such as Shannon’s index (Kers & Saccenti, 2021). Therefore, we split our meta-analysis into two separate analyses for each metric as they provide complementary information about the impacts of plastics on the microbiome, though not all papers provided information on both metrics. We obtained effect sizes by extracting the means and standard deviations for Shannon’s index and relative abundance per taxon from each paper as follows: we either used the published sample sizes, means, and standard deviations (or 95% confidence intervals from which SDs were derived), or we contacted the authors in case these statistics were missing. If authors did not respond, we used WebPlotDigitizer 5.0 [31] to extract the means and standard deviations from the data figures. In this case, we were limited to using bar plots showing the means and standard deviations, e.g. boxplots were excluded. This meant the final number of effect sizes included was n=246 from 13 studies for relative abundance, and n=17 effect sizes from 11 studies for Shannon’s Index, (Fig. 8). There were many more relative abundance effect sizes than Shannon’s Indices because relative abundance was typically estimated for each phylum or species, etc., and each study included relative abundance estimates for multiple phyla or species, depending on the selected taxonomic class of the study. There were, at times, multiple Shannon’s Index effect sizes for a particular study if a study had multiple treatment groups with varying plastic sizes or daily dosages.

We also extracted data on plastic particle size and daily dosage of the exposure, if this information was provided, to test their effects as moderators in our meta-analyses. For the relative abundance data, if there were multiple treatment groups of different dosages, we included the highest dosage treatment group. If the dosages were equivalent and there were multiple experimental groups of different plastic sizes, we used the smallest plastic size treatment group. If there were multiple bacterial taxa ranks analysed, we used the most specific taxa rank (i.e. if genus and species were analysed, we used species) and then re-labelled all taxa at the phyla level to enable including phyla as a moderator variable in our meta-analysis. For plastic size, we took the average value (μm) if a range of plastic sizes were used. For treatment dosage, we used the mg plastics per day scaled to body mass [32]. If body weight was not published, we extracted the average body weight from body weight figures using WebPlotDigitizer for each treatment group. All extracted data, whether used in this meta-analysis or not, can be found in the supplementary information.

We used the escalc function in the metafor package (v4.6-0) [33]to calculate the standardised mean differences between the control and treatment groups expressed as Hedges’ g [34]:

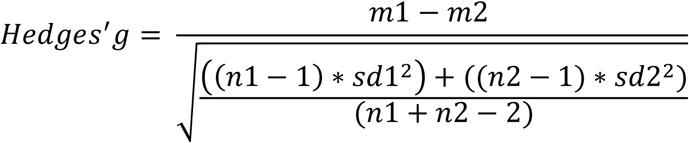

where *m* refers to the mean, *n* refers to the sample size, *sd* refers to the standard deviation, and the numerical labelling refers to the treatment group (1) and the control group (2), i.e. positive effect sizes indicate that microplastic treatment increased gut bacteria metrics.

### Meta-analysis

We used metafor (v4.6-0) [33] in R (v4.4.1) to run multivariate models including study ID and observation ID as random effects [35]. We subsequently added the moderators of plastic size, treatment dosage, and phyla to estimate their respective effects. We used Q-tests to evaluate the heterogeneity among study effect sizes. We tested two different forms of publication biases: bias against publishing small sample size studies without significant results and timing bias against publishing statistically non-significant results. All figures were made using the orchaRd package (v2.0) [35]. See Additional files 2-5 for raw data and meta-analysis R scripts.

## Results

### Effect of microplastics on gut bacteria species alpha diversity

Dietary microplastics did not significantly affect gut bacteria species alpha diversity (0.289, 95% CI -0.362 to 0.941, z-value = 0.871, p = 0.384, Fig. 1). There was significant heterogeneity (Q = 53.728, p < 0.001), indicating substantial variation among effect sizes. We therefore investigated if our moderators, treatment plastic daily dosage and treatment plastic particles size, could explain some of this variation. However, our moderator analysis revealed the daily dosage of plastic particles in the treatment group was not significant (0.941, 95% CI -0.364 to 2.247, z-value = 1.413, p = 0.158, Fig. 2A), nor was the size of plastic particles (0.001, 95% CI -0.011 to 0.013, z-value = 0.216, p = 0.829, Fig. 2B). We found adding these moderators reduced heterogeneity compared to the intercept only model, but the residual heterogeneity remained substantial (QE = 453.446, p < 0.001). One study showed a markedly larger effect size than the rest. Upon closer inspection, the treatment effect appeared similar when comparing the means across treatment groups, however the standard deviations were much smaller compared with the other studies, resulting in a larger Hedges’ g effect size value. We could not identify a specific reason for the much smaller standard deviation in this study. Removing this study from our analysis yielded an indistinguishable overall effect size.

**Figure 1.**
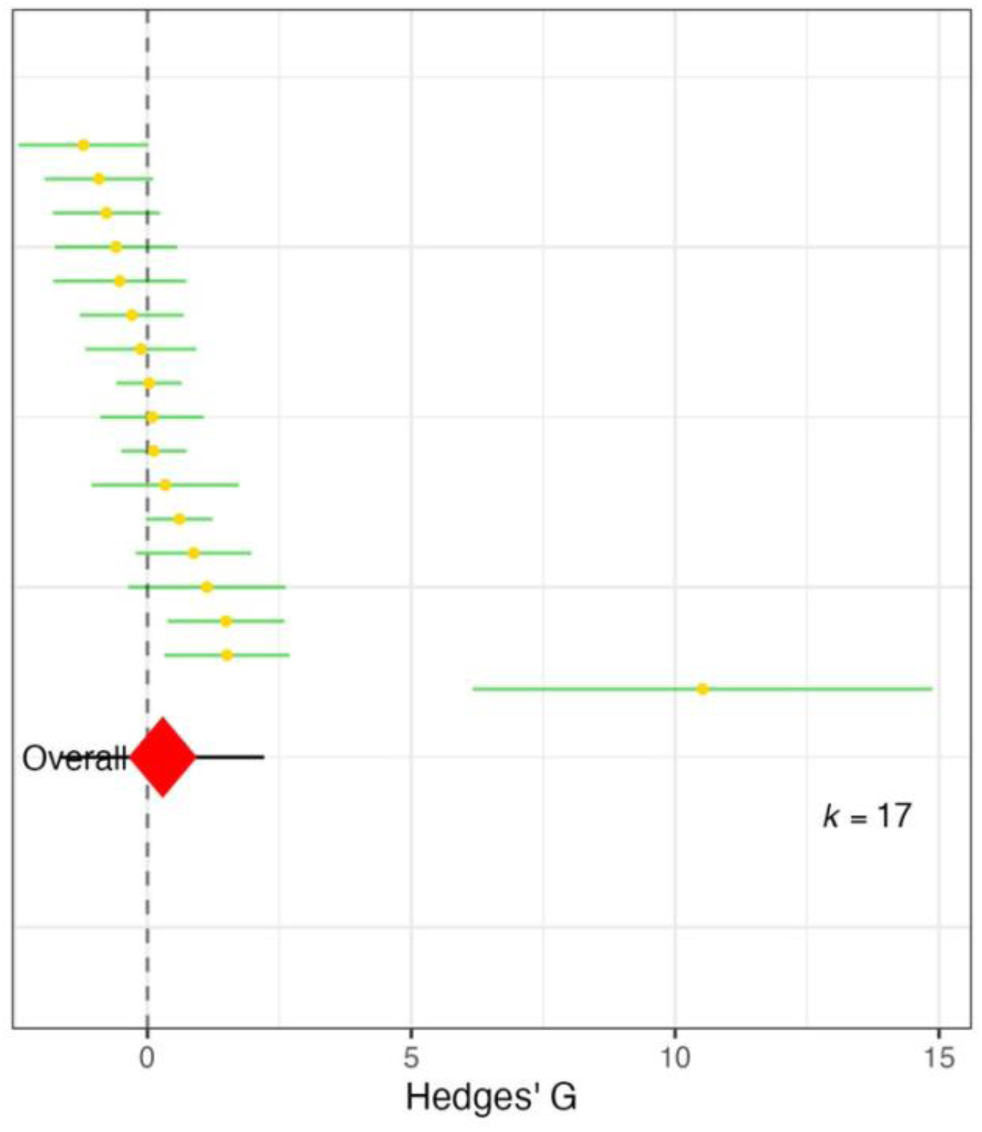
Forest plot of Hedges’ g effect sizes of microplastics impact on gut bacteria alpha diversity. Yellow circles show the meta-analysis effect size point estimate for each set of control vs. treatment means, i.e. positive effect sizes indicate microplastic treatment increased gut bacteria alpha diversity (Shannon index). Horizontal bars indicate 95% confidence intervals. The red diamond indicates the overall effect size estimate where diamond width reflects the predictability interval. *K* denotes the total effect size count.

**Figure 2.**
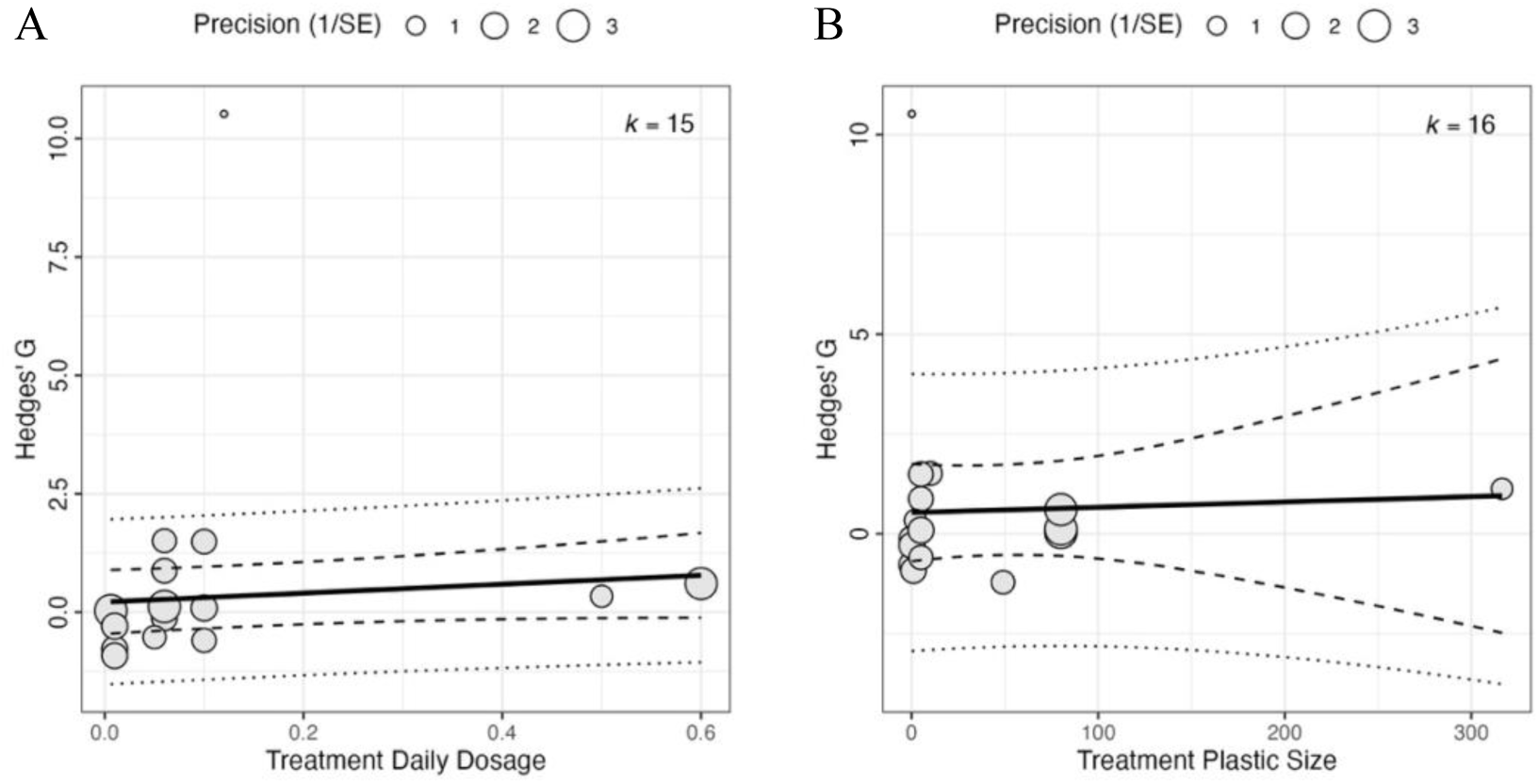
Moderator estimates in the meta-analysis of microplastics impact on gut bacteria species alpha diversity. **(A)** Effect of plastic particles daily dosage on gut bacteria species alpha diversity. **(B)** Effect of particle size on gut bacteria species alpha diversity. The bold lines indicate the moderators’ regression lines. The dashed lines indicate the 95% CI and the dotted lines indicate the predictability intervals. *K* denotes the total number of effect sizes in the moderator analyses. Bubble size reflects the weight of the effect size in the meta-analysis. Gut bacteria alpha diversity is estimated by Shannon diversity index.

We found no evidence for small sample size publication bias (0.721, 95% CI -7.036 to 8.478, z-value = 0.182, p = 0.856, Fig. 3A), or time delay publication bias (0.397, 95% CI - 0.254 to 1.048, z-value = 1.194, p = 0.232, Fig. 3B).

**Figure 3.**
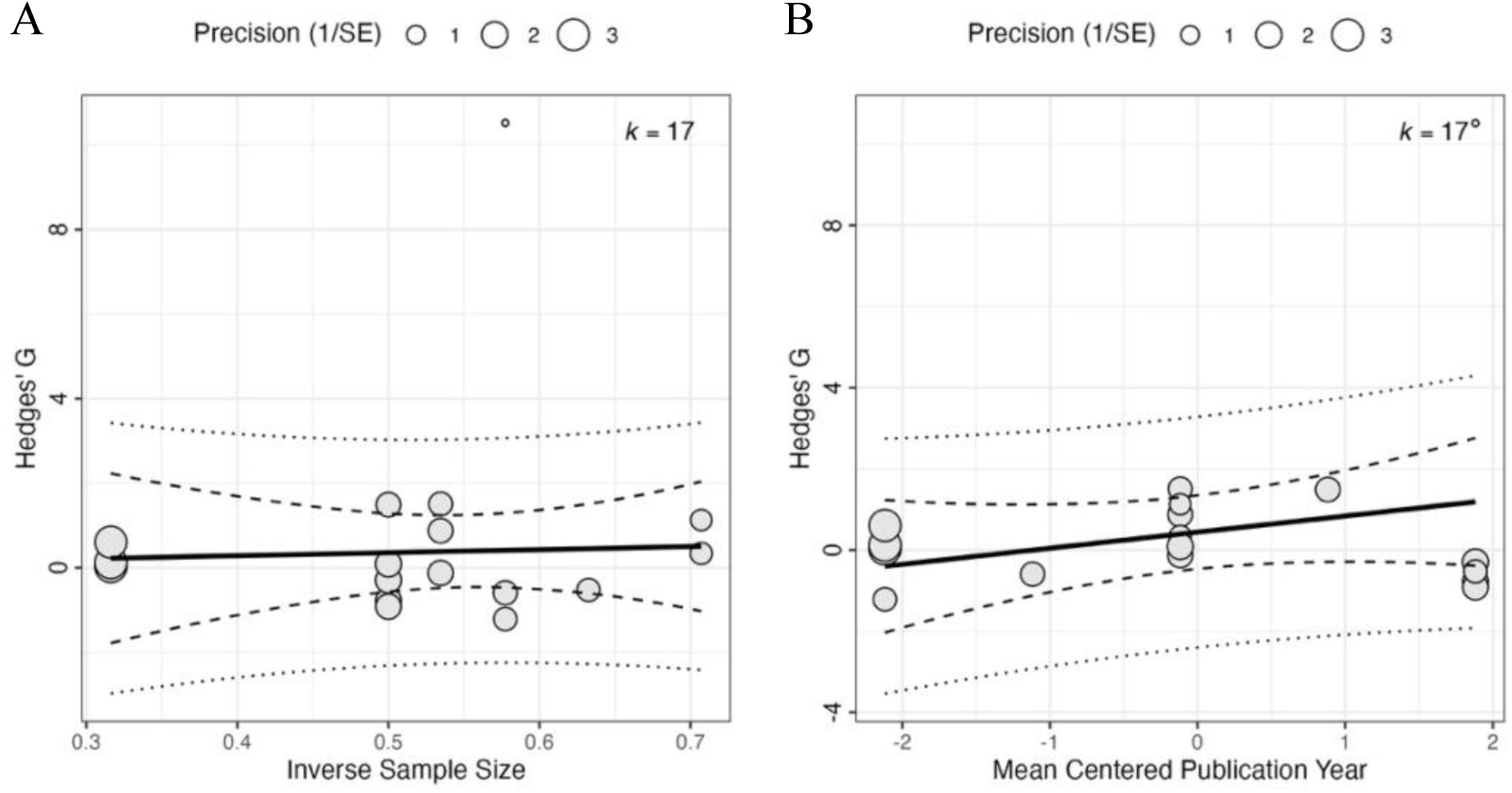
Publication bias for studies on microplastics effect on gut alpha diversity. **(A)** Publication bias against small sample size. **(B)** Publication bias against time delay. The bold line indicates the moderator regression line. The dashed lines indicate the 95% CI and the dotted lines indicate the predictability intervals. *K* denotes the total number of effect sizes in the moderator analysis. Bubble size reflects the weight of the effect size in the meta-analysis. Gut bacteria alpha diversity is estimated by Shannon diversity index.

### Effect of microplastics on the relative abundance of gut bacteria across phyla

We found significant heterogeneity in the effect sizes for relative abundance of gut bacteria across phyla (Q = 714.059, p < 0.001), but the overall effect size was close to zero and not significant (0.503, 95% CI -0.082 to 1.089, z-value = 1.685, p = 0.092, Fig. 4A). However, as the effects of microplastics could either increase or decrease the relative abundance of specific bacterial species, we also looked at the absolute change in relative abundance in response to the treatment using the same data. We then found a significant overall effect size of 1.189 (95% CI 0.704 to 1.674, z-value = 4.805, p < 0.001) suggesting strong evidence for microplastic exposure to induce shifts in the relative abundance of gut bacteria across phyla (Fig. 4B). The negative and positive responses to microplastics treatment shown in Fig. 4A appear to cancel out each other resulting in an overall effect size close to zero. Based on this analysis alone, one could erroneously conclude that microplastics treatment does not affect the gut microbiome composition. Only after looking at the absolute response in Fig 4B, it became clear that there is strong support for microplastics treatment to affect the gut microbiome composition as the relative abundance of bacteria species appears moderately to strongly affected (either negatively or positively) in many cases.

**Figure 4.**
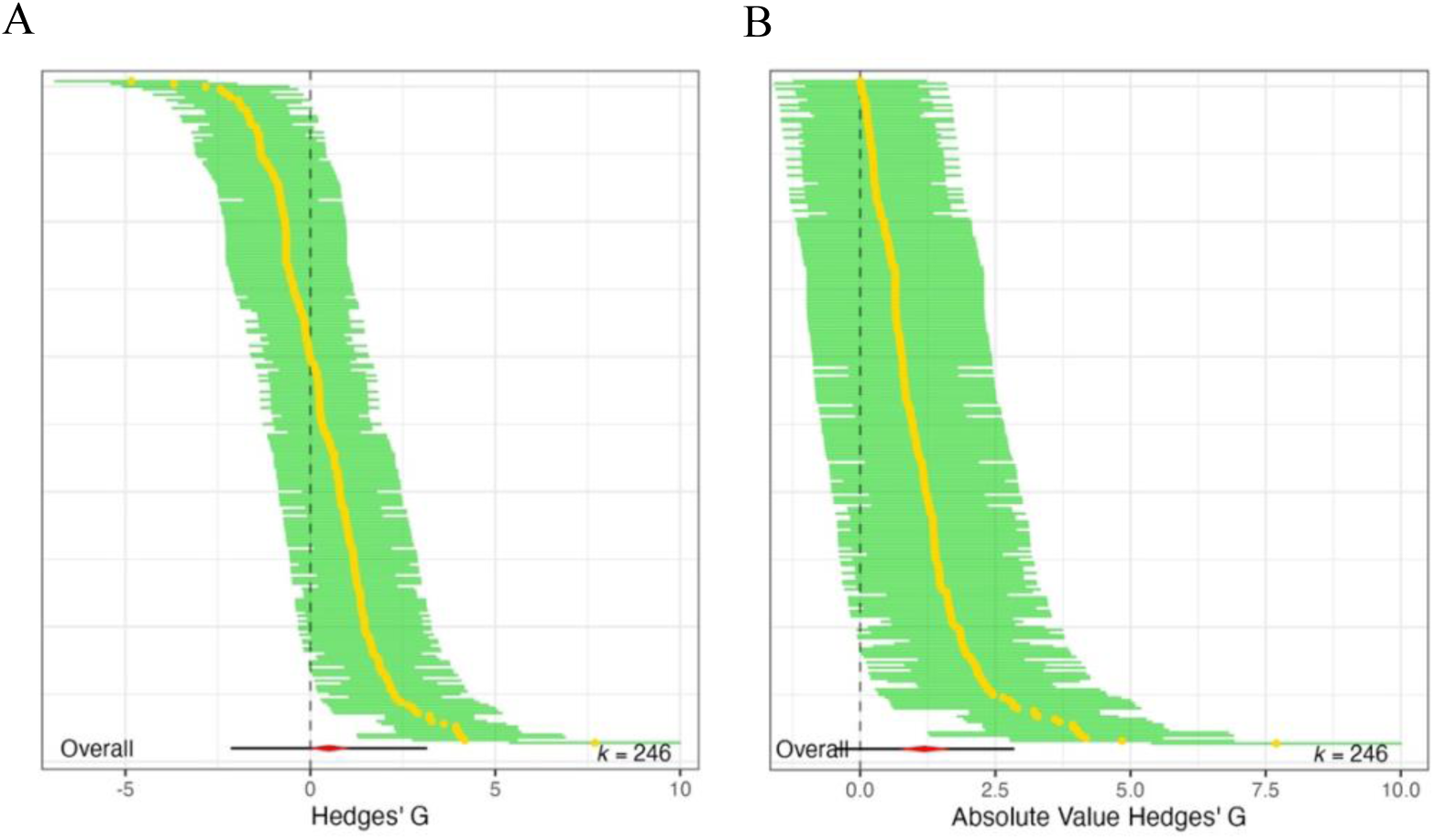
Forest plots of Hedges’ g effect sizes of microplastics impact on gut bacteria relative abundance. (A) Relative abundance across phyla. (B) Absolute changes in relative abundance across phyla, using the same data.Yellow circles show the meta-analysis effect size point estimate for each set of control vs. treatment means, i.e. positive effect sizes indicate microplastic treatment increased the relative abundance of gut bacteria across phyla. Horizontal bars indicate 95% confidence intervals. The red diamond indicates the overall effect size estimate where diamond width reflects the predictability intervals. *K* denotes the total effect size count.

Continuing with the absolute changes iin abundance, our moderator analysis revealed the daily dosage of plastic particles in the treatment group was not significant (-1.695, 95% CI -3.840 to 0.449, z-value = -1.549, p = 0.121, Fig. 5A). On the other hand, the size of plastic particles in the treatment group significantly predicted the absolute response to microplastics exposure (0.042, 95% CI 0.017 to 0.067, z-value = 3.327, p < 0.001, Fig.5B). This finding shows that larger microplastic particles have stronger effects on the relative abundance of gut bacteria across phyla, contrary to what we predicted. While adding these moderators reduced heterogeneity, the residual heterogeneity remained substantial (QE = 166.196, p = 0.961).

**Figure 5.**
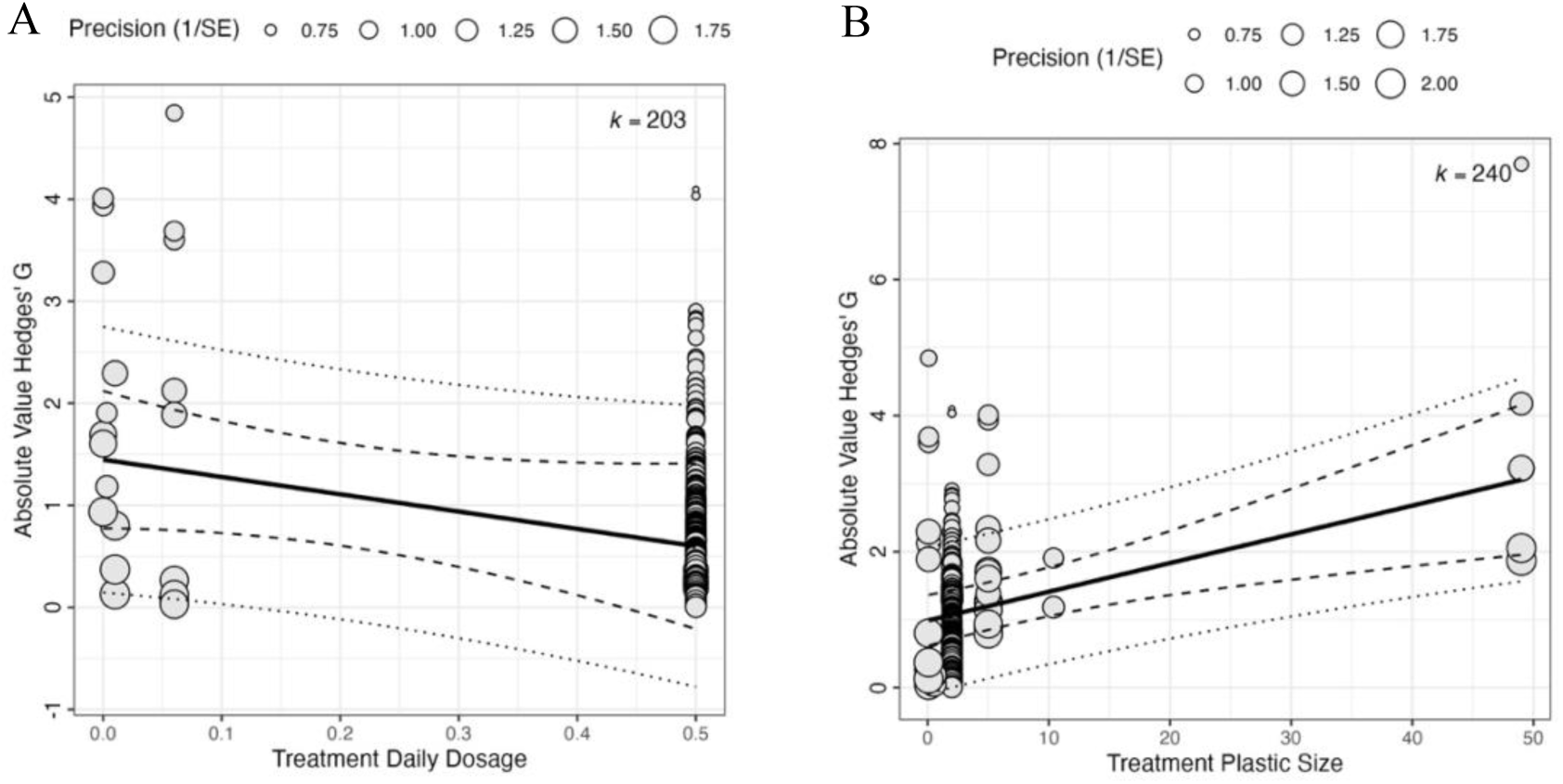
Moderator estimates in the meta-analysis of microplastics impact on relative abundance of gut bacteria. **A)** Effect of plastic particles daily dosage on gut bacteria relative abundance across phyla. **(B)** Effect of particle size on gut bacteria relative abundance across phyla. The bold lines indicate the moderators’ regression lines. The dashed lines indicate the 95% CI and the dotted lines indicate the predictability intervals. *K* denotes the total number of effect sizes in the moderator analyses. Bubble size reflects the weight of the effect size in the meta-analysis.

Returning to relative changes in abundance, the phyla of bacterial species present in the mice gut microbiome was a significant moderator, reducing heterogeneity compared to the intercept only model, though the residual heterogeneity remained substantial (QE = 684.982, p < 0.001). Looking more in detail the microplastics exposure significantly increased relative abundance for two of the 12 phyla, *Bacillota* (0.689, 95% CI 0.037 to 1.341, z-value = 2.072, p = 0.038, Fig. 6) and *Pseudomonadota* (1.097, 95% CI 0.286 to 1.909, z-value = 2.649, p = 0.008, Fig. 6), highlighting the need to consider the possibility of taxa specific responses.

**Figure 6.**
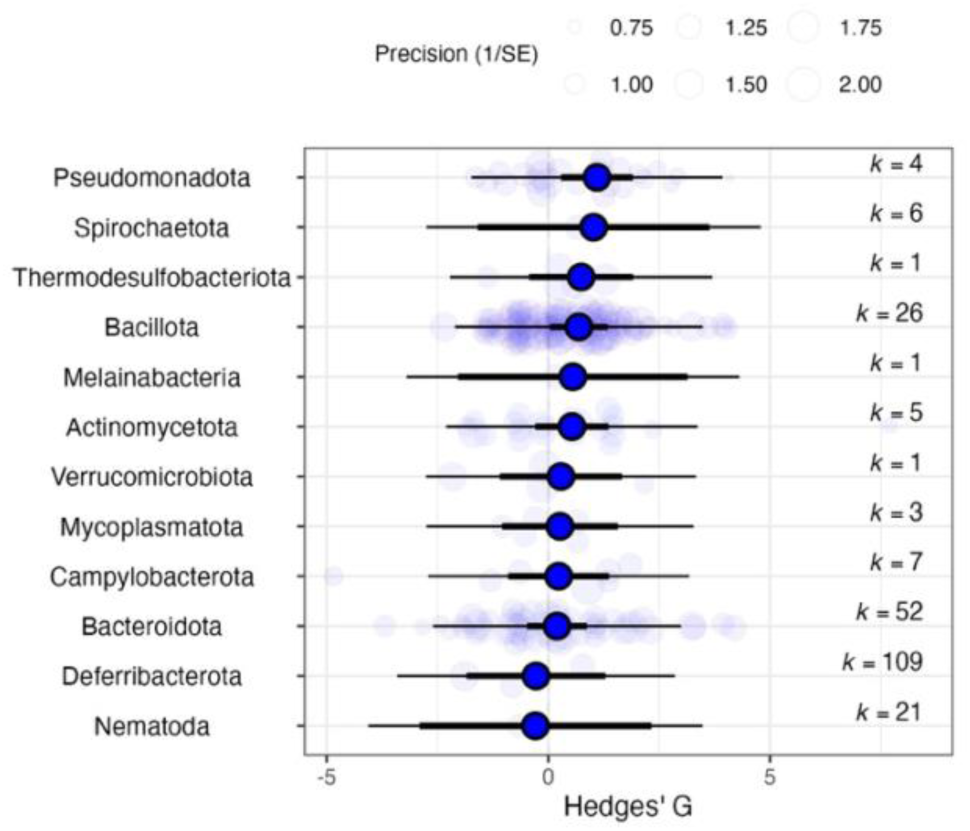
Effect of phyla on the relative abundance for each gut bacteria phyla. The bold lines indicate the 95% CI. *K* denotes the total number of effect sizes in the moderator analyses. Bubble size reflects the weight of the effect size in the meta-analysis.

Finally, we found no evidence for small sample size publication bias (-4.142, 95% CI -8.877 to 0.593, z-value = -1.715, p = 0.086, Fig. 7A), or time delay publication bias (-0.102, 95% CI -0.440 to 0.237, z-value = -0.589, p = 0.556, Fig. 7B).

**Figure 7.**
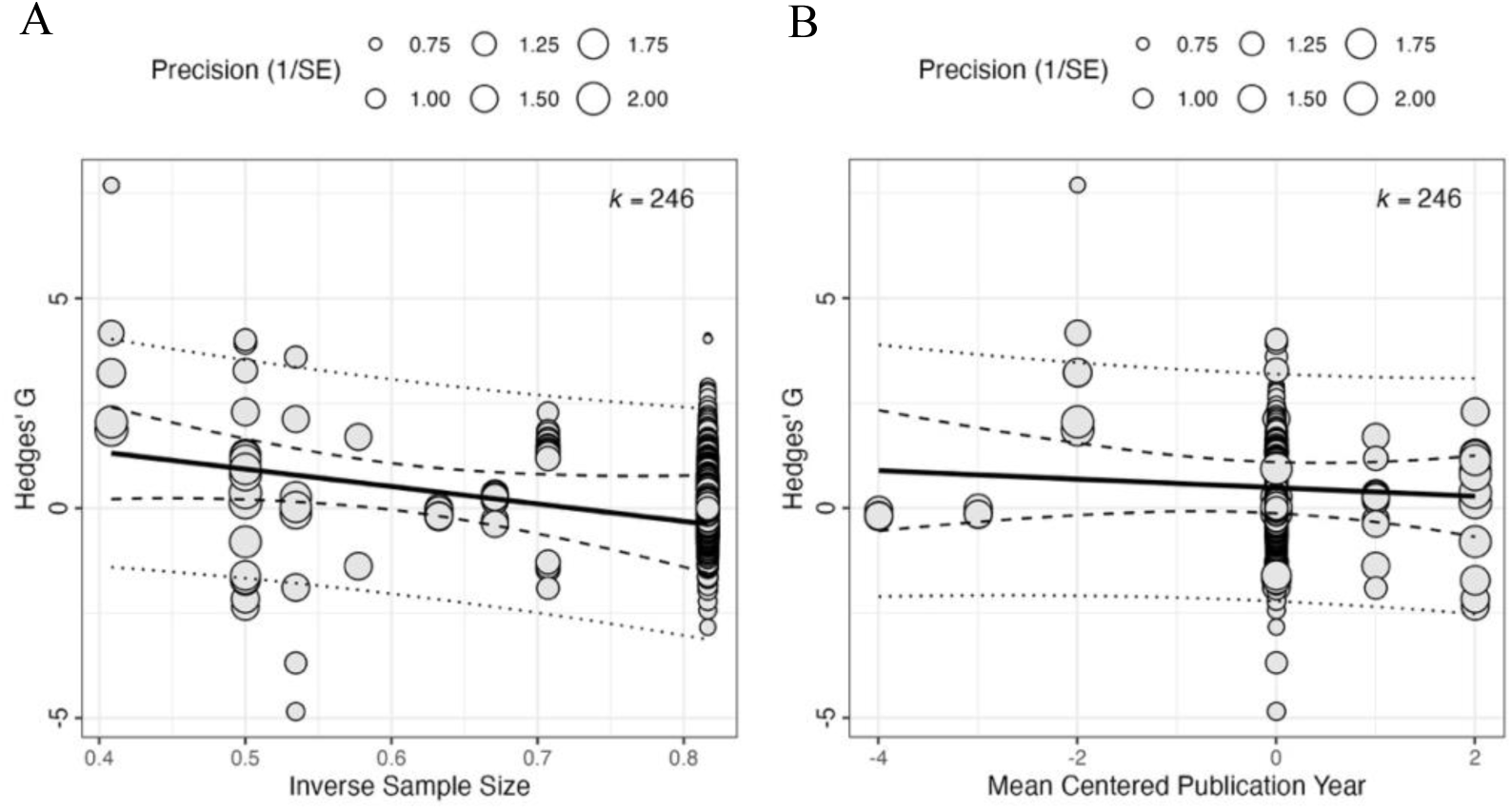
Publication bias for studies on microplastics effect on relative abundance of gut bacteria across phyla. **(A)** Publication bias against small sample size. **(B)** Publication bias against time delay. The bold line indicates the moderator regression trend line. The dashed lines indicate the 95% CI and the dotted lines indicate the predictability intervals. *K* denotes the total number of effect sizes in the moderator analysis. Bubble size reflects the weight of the effect size in the meta-analysis.

**Figure 8.**
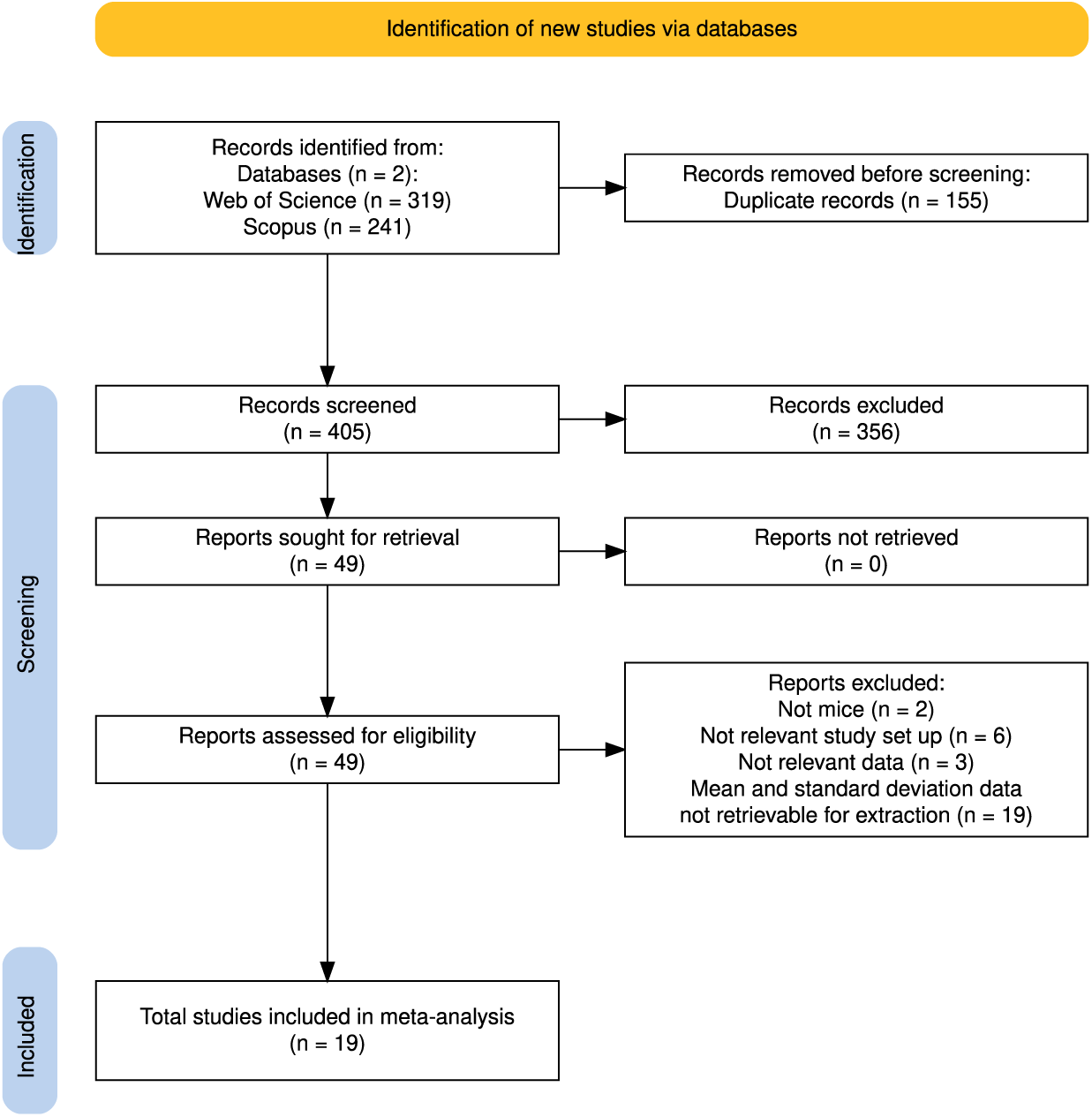
Preferred Reporting Items for Systematic Reviews and Meta-Analyses (PRISMA) diagram of screened studies for meta-analysis. Adapted from [30].

## Discussion

This is the first meta-analysis that tested if experimental dietary exposure to microplastics affects the gut microbiome. Contrary to our initial hypothesis, we did not find a significant overall impact of microplastic exposure on relative abundance of gut bacteria across phyla and Shannon diversity index. However, as microplastics could induce dysbiosis which is typically manifested in a disrupted microbiome balance, the relative abundance of specific species could both decrease or increase. We therefore looked at the absolute change in response to dietary microplastics exposure and then found a significant overall impact of exposure on relative abundance, indicative of heterogeneous effects of microplastics on specific species across phyla. More in-depth meta-analysis revealed two of the twelve bacteria phyla that were investigated (*Bacillota* and *Pseudomonadota*) showed a significant positive response to microplastics exposure (Fig 6). These phyla-specific effects indicate that certain bacterial phyla in the gut microbiome may be relatively more resilient to the influences of the accumulation of microplastics in the gut epithelia, possibly enabling them to thrive and outcompete more vulnerable phyla [8, 26, 27]. Improved phyla level differentiation in future studies will be crucial for improving our understanding of the health consequences of microplastics ingestions as different types of bacteria may have differing functions in the gut microbiome. Our current meta-analysis may therefore not provide the full picture about the detailed interplay between microplastics, microbiome dynamics, and their health implications. Our finding of a significant increase in two specific phyla raises important questions, which we examine in the next paragraphs.

### How comprehensive are the twelve phyla currently included in our analysis, and could there be other important phyla that have not yet been considered?

The twelve included phyla in our meta-analysis represent extractable data from 19 studies. Characterization of the gut microbiome of n=600 lab mice has revealed at least 29 classifiable phyla [36]. The most abundant phyla found in this study were *Bacillota* at 55.75%, *Bacteroidota* at 37.02%, *Pseudomonadota* at 4.05%, *Actinomycetota* at 1.98%, and *Tenericutes* (also known as *Mycoplasmatota*) at 1.09%, with all other phyla at less than 1% [36]. All five of these were reflected in our twelve phyla from extracted data, which would indicate that despite a total of at least 29 classifiable phyla found in lab mice, our picture of twelve phyla is relatively comprehensive as we have already included seven phyla that make up less than 1% of the microbiome of lab mice. Nonetheless, it is important to note that natural mice populations typically have more diverse microbiome compositions compared to lab mice, which represents a limitation of studies that use lab mice [37].

### Does increased relative abundance of Bacillota and Pseudomonadota underpin gut dysbiosis?

Out of the twelve included phyla, the statistically significant increase in the relative abundance of *Bacillota* and *Pseudomonadota* was found to be consistent with existing literature. Specifically, a systematic review found most data show microplastic ingestion is associated with dysbiosis where both *Bacillota* and *Pseudomonadota* increases, however, looking into *Pseudomonadota* further, the subgroups of *Alphaproteobacteria*, *Beta*p*roteobacteria* and *Gammaproteobacteria* have been found to decrease [11]. This difference between *Pseudomonadota* increasing as a phylum, but certain subgroups within it decreasing, points to the fact that while phyla-level differentiation is stronger than a blanket diversity index, a phyla is still a broad taxonomic category and further differentiating by more precise taxonomic categories might be needed to derive full understanding of how the microbiome is reacting to microplastic exposure, which some studies have done and are discussed in the next question. Deeper investigation into the functions of the two phyla that significantly increased in relative abundance tells us *Bacillota* is important for carbohydrate metabolism in the gut microbiome of humans and is often most dominant by composition in humans and mice, whereas *Pseudomonadota* has been found to include facultative anaerobes which are often opportunistic pathogens [38]. Thus, while statistically significant changes in these two phyla are indeed indicative of dysbiosis, *Pseudomonadota* may have specific negative implications for healthy microbiome functioning.

### Do specific bacteria species drive the increased relative abundance of Bacillota and Pseudomonadota?

While there is some variation in specific families, genera, or species reported, the common patterns found across studies include an increase in species from the family *Muribaculaceae*, genus *Lactobacillus*, and genus *Ruminiclostridium* in response to microplastic exposure in mice [39–41]. These patterns are consistent with our analysis as *Muribaculaceae* is from the phyla *Bacteroidota*, which showed a non-significant increase in relative abundance in our meta-analysis, and *Lactobacillus* and *Ruminiclostridium* are both from the phyla *Bacillota,* which showed a significant increase in relative abundance. *Muribaculaceae* is associated with regulation of gut and immune function [42], *Lactobacillus* is associated with carbohydrate metabolism and pathogen protection [43], and *Ruminiclostridium* is associated with cellulolytic activity and obesity regulation [44].

### Can microplastic-induced changes in the microbiome be linked to medical conditions such as chronic inflammation?

Increases in *Pseudomonadota* are linked with elevated oxidative stress in the intestinal epithelial cells, associated with inflammation [3]. It has been suggested that its increase in relative abundance could reflect a competitive advantage as bacteria in this phyla are relatively aerotolerant and therefore more resistant against oxidative stress [26]. Within *Pseudomonadota*, the family *Enterobacteriaceae* has been shown to quickly increase in relative abundance with oxidative stress and inflammation [45]. Whether such changes are symptomatic, or possibly functionally linked with gut health remains unclear. Future studies can incorporate germ free mice to allow for more meaningful studies of causation vs. correlation with inflammation [26] as well as focus on comparing patterns of relative abundance per bacterial species associated with inflammation such as *Enterobacteriaceae*.

### To what extent are lab dosages of microplastics realistic and representable for organisms in the natural environment?

Our meta-analysis supports the existence of experimental effects of microplastics ingestions on the gut microbiome, in controlled laboratory conditions. Whether our findings are relevant for wild animal populations remains to be seen. A limitation of previous captive experimental studies is that they typically use one polymer whereas in the wild, it is likely that animals are exposed to a cocktail of different types of polymers such as polyethylene, polypropylene, and polystyrene [6, 8, 16]. In addition, captive studies usually use sterile particles, whereas in the wild, microplastics can occur as biofilm [8] or be coated with pesticides and other toxic chemicals [41]. A key question therefore remains whether there could be additive or multiplicative effects of these pollution factors. To begin addressing such questions we need experimental studies on outbred, or non-model organisms using realistic doses of microplastics. For example, a recent study investigated the effects of naturally occurring doses of microplastics on growth, oxidative stress and gut telomere length in African clawed frogs and this study found no significant treatment effects on oxidative stress and gut epithelial telomere length [46]. More studies like these will be essential for improving our understanding of the health consequences of environmental microplastics pollution. Finally, it will be particularly important to increase our understanding of such effects in terrestrial populations as the current focus has largely been on the impacts of microplastics on species in marine environments [2, 8, 13].

## Conclusions

Given existing studies on the effects of microplastic exposure on the microbiome demonstrated contrasting findings, this meta-analysis was performed to test if experimental dietary exposure to microplastics affects the gut microbiome. Our results demonstrated support for a significant impact of microplastic exposure on the relative abundance of gut bacteria, and specifically, for an increase in the relative abundance of certain gut bacterial phyla such as *Bacillota* and *Pseudomonadota.* Deeper analysis into these phyla tell us that subgroups within *Pseudomonadota,* for example, have shown contrasting directional impacts from microplastics, [11], emphasizing future studies may further differentiate by more precise taxonomic categories beyond phyla such as species for more discrete patterns of impact to be studies. Future studies may also include non-model organisms with a mix of plastic polymer types for greater translation to wild populations [6, 8, 16], germ free mice for more meaningful differentiation of causation vs. correlation with inflammation [26], or more terrestrial organisms as they are relatively understudied today.

## Supporting information

Additional file 2- Shannon index data

Additional file 3- Shannon index meta-analysis

Additional file 4- Relative abundance data

Additional file 5- Relative abundance meta-analysis

## Declarations

### Ethics approval and consent to participate

Not applicable

### Consent for publication

Not applicable

### Availability of data and material

All data generated or analyzed during this study are included in this published article (and its supplementary information files).

### Authors’ contributions

This study was conceived by JB and DD. AK performed the literature search, extracted the relevant statistics for effect sizes, performed the meta-analyses, and wrote the first draft of the manuscript. All authors contributed to revisions.

## Acknowledgements

In alphabetical order, thank you to Liehai Hu and Hengyi Xu for coordinating and/or sharing requested data used in this meta-analysis. Thank you to Rachel Reid for aiding in the meta-analysis code. Thank you to Adam Dobson for feedback on the manuscript draft.

## Additional files

Additional file 1- Table S1.xlsx includes Table S1. Studies Included in this meta-analysis (n=19). Table S1 lists out the treatment daily dosage (mg/day), average treatment plastic size (μm), and available phyla for the 19 studies whose data was included in this meta-analysis.

Additional file 2- Shannon index data.csv includes the raw data for the first meta-analysis on gut bacteria species alpha diversity as measured by Shannon index.

Additional file 3- Shannon index meta-analysis.Rmd includes the R scripts for the first meta-analysis on gut bacteria species alpha diversity as measured by Shannon index.

Additional file 4- Relative abundance data.csv includes the raw data for the second meta-analysis on the relative abundance of gut bacteria across phyla.

Additional file 5- Relative abundance meta-analysis.Rmd includes the R scripts for the second meta-analysis on the relative abundance of gut bacteria across phyla.

