## Additional file 5- Relative abundance meta-analysis for "The impact of microplastics on the mice gut microbiome: a meta-analysis": Additional file 5- Relative abundance meta-analysis.Rmd

````
---
title: "Step 1 to Step 4 Relative Abundance Meta-Analysis Manuscript Final"
author: "Anika Kalra and Rachel Reid"
date: "July 2024"
output:
  html_document:
    df_print: paged
---
*Script Aim: Go through 4 steps to conduct meta-analysis for relative abundance of gut microbiome bacteria*

**Step 1: Calculating hedges g using escalc function in metafor R package**
```{r}
#Install and load packages
library(pacman)
pacman::p_load(dplyr,ggplot2,meta,metafor, robvis, ape,rotl,Hmisc, circlize,
               dendextend, tidyverse, gridExtra, phytools, plyr, boot, brms)

#Read in your own dataset
Data <- (read_csv("Additional file 4- Relative abundance data.csv"))
Data$TreatmentDailyDosage<-as.numeric(Data$TreatmentDailyDosage)
Data$TreatmentPlasticSize<-as.numeric(Data$TreatmentPlasticSize)
```

#Calculating standardized mean differences using the escalc function
```{r}
#n1i and n2i correspond to sample sizes for group 1 and 2, m1i and m2i correspond to means for groups
#1 and 2, sd1i and sd2i correspond to standard deviation for groups 1 and 2
SMD<-escalc(measure = "SMD", n1i = Data$TreatmentSampleSize, n2i=Data$ControlSampleSize,
             m1i=Data$TreatmentMean, m2i=Data$ControlMean, sd1i=Data$TreatmentSD, 
             sd2i=Data$ControlSD)

#Combining effect size columns to your dataset
Data_new<-bind_cols(Data,SMD)

#add in absvalue of effect sizes
Data_new<- Data_new%>%
  mutate(yiabsval= abs(yi))

#Saving your dataset with the effect size columns now included
Data_final<- as.data.frame(Data_new)
write.csv(Data_final, "Data_final_Abundance.csv")
```

**Step 2: Running Meta-analysis models using rma.mv function in metafor R package**
```{r}
#Install and load packages
pacman::p_load(dplyr,ggplot2,meta,metafor, robvis, ape,rotl,Hmisc, circlize,
               dendextend, tidyverse, gridExtra, phytools, plyr, boot, brms,
               ggpubr, metaviz, orchaRd, R.rsp, patchwork)
pacman::p_load(devtools, tidyverse, metafor, patchwork, R.rsp, emmeans)

devtools::install_github("daniel1noble/orchaRd", force = TRUE)
```

#Intercept only model
```{r}
#yi corresponds to hedges g, v is your variance, obs_ID is your effect size level ID
Obs_ID <- c(1,2,3,4,5,6,7,8,9,10,11,12,13,14,15,16,17,18,19,20,21,22,23,24,25,26,27,28,29,30,31,32,33,34,35,36,37,38,39,40,41,42,43,44,45,46,47,48,49,50,51,52,53,54,55,56,57,58,59,60,61,62,63,64,65,66,67,68,69,70,71,72,73,74,75,76,77,78,79,80,81,82,83,84,85,86,87,88,89,90,91,92,93,94,95,96,97,98,99,100,101,102,103,104,105,106,107,108,109,110,111,112,113,114,115,116,117,118,119,120,121,122,123,124,125,126,127,128,129,130,131,132,133,134,135,136,137,138,139,140,141,142,143,144,145,146,147,148,149,150,151,152,153,154,155,156,157,158,159,160,161,162,163,164,165,166,167,168,169,170,171,172,173,174,175,176,177,178,179,180,181,182,183,184,185,186,187,188,189,190,191,192,193,194,195,196,197,198,199,200,201,202,203,204,205,206,207,208,209,210,211,212,213,214,215,216,217,218,219,220,221,222,223,224,225,226,227,228,229,230,231,232,233,234,235,236,237,238,239,240,241,242,243,244,245,246,247,248,249,250,251,252,253,254,255,256,257,258,259,260,261,262)
Data_final$Obs_ID <- Obs_ID

#check that TreatmentDailyDosage and TreatmentPlasticSize are numeric
glimpse(Data_final)

model1 <- rma.mv(yi = yi, 
                 V = vi, 
                 mods = ~ 1, 
                 random = list(~1|Study,
                               ~1|Obs_ID),
                 data=Data_final, 
                 method="REML")

summary(model1)
```
```{r}
#Model 1 with absval effect sizes
model1absval <- rma.mv(yi = yiabsval, 
                 V = vi, 
                 mods = ~ 1, 
                 random = list(~1|Study,
                               ~1|Obs_ID),
                 data=Data_final, 
                 method="REML")

summary(model1absval)
```

#Add in Moderators
```{r}
#Model 2 added moderator, TreatmentDailyDosage of effect sizes
model2 <- rma.mv(yi = yi, 
                 V = vi,
                 mods = ~ TreatmentDailyDosage,  
                 random = list(~1|Study,
                               ~1|Obs_ID),
                 data=Data_final, 
                 method="REML")
summary(model2)
```

```{r}
#Model 2 added moderator, TreatmentDailyDosage of absval effect sizes
model2absval <- rma.mv(yi = yiabsval, 
                 V = vi,
                 mods = ~ TreatmentDailyDosage,  
                 random = list(~1|Study,
                               ~1|Obs_ID),
                 data=Data_final, 
                 method="REML")
summary(model2absval)
```
```{r}
#Model 3 added another moderator, TreatmentPlasticSize of effect sizes
model3 <- rma.mv(yi = yi, 
                 V = vi,
                 mods = ~ TreatmentPlasticSize,  
                 random = list(~1|Study,
                               ~1|Obs_ID),
                 data=Data_final, 
                 method="REML")
summary(model3)
```

```{r}
#Model 3 added another moderator, TreatmentPlasticSize of absval effect sizes
model3absval <- rma.mv(yi = yiabsval, 
                 V = vi,
                 mods = ~ TreatmentPlasticSize,  
                 random = list(~1|Study,
                               ~1|Obs_ID),
                 data=Data_final, 
                 method="REML")
summary(model3absval)
```

```{r}
#Model 4 last moderator, phylum alone 
model4 <- rma.mv(yi = yi, 
                 V = vi,
                 mods = ~ Phylum-1,  
                 random = list(~1|Study,
                               ~1|Obs_ID),
                 data=Data_final, 
                 method="REML")
summary(model4)
```
```{r}
#Model 5 added both moderators, TreatmentDailyDosage and TreatmentPlasticSize of absval effect sizes
model5absval <- rma.mv(yi = yiabsval, 
                 V = vi,
                 mods = ~ TreatmentDailyDosage
                 + TreatmentPlasticSize,  
                 random = list(~1|Study,
                               ~1|Obs_ID),
                 data=Data_final, 
                 method="REML")
summary(model5absval)
```

**Step 3: Calculating publication bias using rma.mv function in metafor R package**
```{r}
#Install and load packages
pacman::p_load(dplyr,ggplot2,meta,metafor, robvis, ape,rotl,Hmisc, circlize,
               dendextend, tidyverse, gridExtra, phytools, plyr, boot, brms,
               ggpubr, metaviz, orchaRd, R.rsp, patchwork)
pacman::p_load(devtools, tidyverse, metafor, patchwork, R.rsp, emmeans)

#Creating dataframe to look at publication bias
df_pb <- Data_final %>% ##using your original dataset 
  filter(!is.na(yi)) %>% 
  filter(!is.na(vi)) %>% 
  mutate(s_weights = 1/vi,
         sqrt_inv_eff_ss = sqrt((1/ControlSampleSize) + (1/TreatmentSampleSize)),#inverse sample size, this will be used to calculate small study effects
         inv_eff_ss = (1/ControlSampleSize) + (1/TreatmentSampleSize),
         publication_year_c = PublicationYear - mean(PublicationYear))#mean centered pub year will be used to calculate timelag bias
```

#Small-study effects publication bias
```{r}
model_pub_bias1 <- rma.mv(yi = yi, 
                          V = vi, 
                          mods = ~ sqrt_inv_eff_ss, 
                          random = list(~1|Study, 
                                        ~1|Obs_ID),
                          data=df_pb, 
                          method="REML")
summary(model_pub_bias1)

#Calculating R squared To look at if there is small study bias
r2_ml(model_pub_bias1)

#Make plot
bubblepubbias1<-orchaRd::bubble_plot(model_pub_bias1, group = "Study", mod = "sqrt_inv_eff_ss",
                     legend.pos = "top.out", est.col="black", cb=TRUE, alpha=5, 
                     xlab= "Inverse Sample Size", ylab= "Hedges' G")

#Saving the plot
ggsave(filename = "Final_SmallStudyEffectsAbundance.jpeg", 
       plot = bubblepubbias1, 
       device = "jpeg", 
       width = 110, 
       height = 120, 
       units = "mm")
```

#Time lag publication bias
```{r}
model_pub_bias2 <- rma.mv(yi = yi, 
                          V = vi, 
                          mods = ~ publication_year_c, 
                          random = list(~1|Study, 
                                        ~1|Obs_ID),
                          data=df_pb, 
                          method="REML")
summary(model_pub_bias2)

#Calculating R squared to look at timelag bias
r2_ml(model_pub_bias2)

#Make plot
bubblepubbias2<-orchaRd::bubble_plot(model_pub_bias2, group = "Study", mod = "publication_year_c",
                                     legend.pos = "top.out", est.col="black", cb=TRUE, alpha=5, 
                                     xlab= "Mean Centered Publication Year", ylab= "Hedges' G")

#Saving the plot
ggsave(filename = "Final_TimeLagBiasAbundance.jpeg", 
       plot = bubblepubbias2, 
       device = "jpeg", 
       width = 110, 
       height = 120, 
       units = "mm")
```

**Step 4: Plotting your data using orchard_plot,bubble_plot, caterpillars functions in orchaRd package**
```{r}
#Install and load packages
library(orchaRd)
```

#Plotting categorical moderator data, Phylum, using orchard_plot
```{r}
Base_plot4<-orchard_plot(object = model4, ##The corresponding model you are wanting to plot
                         mod = "Phylum", #This is what will appear on your y axis 
                         group= "Study",
                         cb =TRUE, #If TRUE uses colour blind friendly colours
                         g=FALSE, #If true displays number of grouping levels for each level of mod, in this case it would be number of studies
                         angle=360, #the angle of y labels
                         xlab = "Hedges' G",
                         legend.pos = "top.out",
                         alpha=0.05, 
                         trunk.size=1, #size of the mean or central point
                         transfm = "none",
                         k=TRUE)# if true displays number of effect sizes 

#Saving the plot
ggsave(filename = "Final_PhylumAbundance.jpeg", 
       plot = Base_plot4+
scale_x_discrete(limits = sub(x = names(sort(coefficients(model4))),
                                pattern = 'Phylum',
                                replacement = '')) +
  scale_fill_manual(values = rep('Blue', 12)),
       device = "jpeg", 
       width = 140, 
       height = 120, 
       units = "mm")
```

#Plotting continuous moderator data, TreatmentDailyDosage, using bubble_plot 
```{r}
lim_bubble2 <- orchaRd::mod_results(model2absval, mod = "TreatmentDailyDosage", group = "Study")

Bubble2 <-orchaRd::bubble_plot(lim_bubble2, group = "Study", mod = "TreatmentDailyDosage",
                               legend.pos = "top.out", est.col="black", cb=TRUE, alpha=5, 
                               xlab= "Treatment Daily Dosage", ylab= "Absolute Value Hedges' G")

#Saving the plot
ggsave(filename = "Final_TreatmentDailyDosageAbundance.jpeg", 
       plot = Bubble2, 
       device = "jpeg", 
       width = 120, 
       height = 130, 
       units = "mm")
```

#Plotting continuous moderator data, TreatmentPlasticSize, using bubble_plot 
```{r}
lim_bubble3 <- orchaRd::mod_results(model3absval, mod = "TreatmentPlasticSize", group = "Study")

Bubble3 <-orchaRd::bubble_plot(lim_bubble3, group = "Study", mod = "TreatmentPlasticSize",
                             legend.pos = "top.out", est.col="black", cb=TRUE, alpha=5,
                             xlab= "Treatment Plastic Size", ylab= "Absolute Value Hedges' G")

#Saving the plot
ggsave(filename = "Final_TreatmentPlasticSizeAbundance.jpeg", 
       plot = Bubble3, 
       device = "jpeg", 
       width = 120, 
       height = 130, 
       units = "mm")
```

#Plotting intercept only model, using caterpillars
```{r}
Base_plot1<-caterpillars(object = model1, ##The corresponding model you are wanting to plot
                         mod = "1", #This is what will appear on your y axis 
                         group= "Study",
                         g=FALSE, #If true displays number of grouping levels for each level of mod, in this case it would be number of studies
                         xlab = "Hedges' G",
                         transfm = "none",
                         k=TRUE)# if true displays number of effect sizes 

#Saving the plot
ggsave(filename = "Final_InterceptOnlyModelAbundance.jpeg", 
       plot = Base_plot1, 
       device = "jpeg", 
       width = 110, 
       height = 120, 
       units = "mm")

Base_plot1absval<-caterpillars(object = model1absval, ##The corresponding model you are wanting to plot
                         mod = "1", #This is what will appear on your y axis 
                         group= "Study",
                         g=FALSE, #If true displays number of grouping levels for each level of mod, in this case it would be number of studies
                         xlab = "Absolute Value Hedges' G",
                         transfm = "none",
                         k=TRUE)# if true displays number of effect sizes 

#Saving the plot
ggsave(filename = "Final_InterceptOnlyModelAbsValAbundance.jpeg", 
       plot = Base_plot1absval, 
       device = "jpeg", 
       width = 110, 
       height = 120, 
       units = "mm")
```
````
